# Neural correlates of confidence during decision formation in a perceptual judgment task

**DOI:** 10.1101/2023.08.13.553156

**Authors:** Yiu Hong Ko, Andong Zhou, Eva Niessen, Jutta Stahl, Peter H. Weiss, Robert Hester, Stefan Bode, Daniel Feuerriegel

## Abstract

When we make a decision, we also estimate the probability that our choice is correct or accurate. This probability estimate is termed our degree of decision confidence. Recent work has reported event-related potential (ERP) correlates of confidence both during decision formation (the centro-parietal positivity component; CPP) and after a decision has been made (the error positivity component; Pe). However, there are several measurement confounds that complicate the interpretation of these findings. More recent studies that overcome these issues have so far produced conflicting results. To better characterise the ERP correlates of confidence we presented participants with a comparative brightness judgment task while recording electroencephalography. Participants judged which of two flickering squares (varying in luminance over time) was brighter on average. Participants then gave confidence ratings ranging from “surely incorrect” to “surely correct”. To elicit a range of confidence ratings we manipulated both the mean luminance difference between the brighter and darker squares (relative evidence) and the overall luminance of both squares (absolute evidence). We found larger CPP amplitudes in trials with higher confidence ratings. This association was not simply a by-product of differences in relative evidence (which covaries with confidence) across trials. We did not identify postdecisional ERP correlates of confidence, except when they were artificially produced by pre-response ERP baselines. These results provide further evidence for neural correlates of processes that inform confidence judgments during decision formation.

## 1. Introduction

When we make decisions, we also estimate the probability that our choice is accurate or will lead to desired outcomes. These (often implicit) confidence judgments have been conceptualised as ‘second-order’ decisions across a continuous dimension (ranging from being certain that the decision was incorrect, to being certain that the decision was correct), signalling the prospect that a corresponding ‘first-order’ decision is accurate (Yeung & Summerfield, 2012; Pouget et al., 2016; Fleming & Daw, 2017). We can use our sense of confidence as a proxy for objective choice accuracy to rapidly correct errors (Rabbitt & Vyass, 1981; Yeung & Summerfield, 2012) and determine whether we should adjust our decision-making strategies to improve performance (Vickers, 1979; van den Berg et al., 2016a; Desender et al., 2019a).

Two broad classes of models have been proposed to account for confidence judgments in perceptual decision tasks. The first class of‘decisional-locus’ models (as labelled by Yeung & Summerfield, 2012) specify that confidence judgments are (primarily or exclusively) based on information related to features of the first-order decision (e.g., Vickers, 1979; Kiani & Shadlen, 2009; Kiani et al., 2014). This includes a subset of racing accumulator models that predict confidence as a function of the relative extent of accumulated evidence in favour of each choice alternative (e.g., Vickers, 1979; Vickers & Packer, 1982; Ratcliff & Stams, 2009). By contrast, postdecisional locus models describe processes that occur after the time of the first-order decision, such as continued evidence accumulation (Rabbitt & Vyass, 1981; Pleskac & Busemeyer, 2010; Moran et al., 2015; van den Berg et al., 2016b; Desender et al., 2021a; Maniscalco et al., 2021) or computations based on other sources of information (e.g., exerted motoric effort, Fleming & Daw, 2017; Gajdos et al., 2019; Turner et al., 2021a; Overhoff et al., 2022).

Each class of models specifies critical processes that occur over different time windows relative to the first-order decision. Accordingly, researchers have tested for neural correlates of confidence during decision formation and postdecisional time windows using high temporal resolution neural recordings such as electroencephalography (EEG) and analyses of event-related potential (ERP) components. This has produced two distinct sets of findings: one related to the centro-parietal positivity (CPP) component during decision formation (O’Connell et al., 2012; Twomey et al., 2015) and another linked to the postdecisional error positivity (Pe) component (Falkenstein et al., 1991, for reviews see Rausch et al., 2020; Feuerriegel et al., 2022).

### 1.1. ERP Correlates of Confidence During Decision Formation

The CPP component (analogous to the P3b, Twomey et al., 2015) has a morphology that closely resembles accumulation-to-boundary trajectories of decision variables in evidence accumulation models (Ratcliff, 1978; Ratcliff et al., 2016; O’Connell et al., 2012; Kelly & O’Connell, 2013; Twomey et al., 2015). The CPP has therefore been conceptualised as a neural correlate of evidence accumulation trajectories in decision-making tasks. Accordingly, the amplitude of this component at the time of the decision (e.g., immediately preceding a keypress response) has been interpreted as indexing the extent of evidence accumulated in favour of the chosen option (e.g., Philiastides et al., 2014; Gherman & Philiastides, 2018; Steinemann et al., 2018; von Lautz et al., 2019; Feuerriegel et al., 2021; Kelly et al., 2021). Larger stimulus-locked CPP amplitudes have been reported to co-occur with higher confidence ratings (Squires et al., 1973; Gherman & Philiastides, 2015, 2018; Herding et al., 2019; Zakrzewski et al., 2019; Rausch et al., 2020) and also higher stimulus visibility ratings as measured using a perceptual awareness scale (Tagliabue et al., 2019). Larger pre-response amplitudes have also been reported for higher model-estimated (Philiastides et al., 2014) and participant-reported confidence ratings (Grogan et al., 2023; but see Feuerriegel et al., 2022). This has been taken as support for the ‘balance of evidence’ hypothesis proposed in racing accumulator models (e.g., Vickers, 1979; Vickers & Packer, 1982; Smith & Vickers, 1988; Ratcliff & Stams, 2009; discussed in Smith et al, 2022). In these models, a first-order decision is made when one of multiple racing accumulators reaches a decision boundary. Differences in the relative extent of evidence accumulated for chosen and unchosen options at the time of the first-order decision (assumed to be indexed by CPP amplitudes) determine one’s degree of confidence. Here, we note that there are several other decisional-locus models that do not propose the balance of evidence hypothesis (reviewed in Pleskac & Busemeyer, 2010; Yeung & Summerfield, 2012; Fleming & Daw, 2017). These models could also be formulated in ways that specify links to CPP amplitudes during decision formation.

There are, however, several conceptual and measurement issues that complicate the interpretation of these findings. The first issue is that racing accumulator models specify differences in evidence accumulation across choice options *at the time of the decision* (approximated by the time of the motor response). However, the majority of studies reporting positive associations between CPP amplitudes and decision confidence have analysed stimulus-locked ERPs (e.g., Squires et al., 1973; Gherman & Philiastides, 2015, 2018; Herding et al., 2019; Zarkewski et al., 2019; Rausch et al., 2020). Because response times (RTs) vary widely across trials, it is unclear whether these stimulus-locked measures capture amplitude variations that are most pronounced at the time of decision formation. In addition, higher confidence ratings typically co-occur with faster RTs in most perceptual decision tasks (e.g., Johnson, 1939; Vickers & Packer, 1982; Kiani et al., 2014; Rahnev et al., 2020). As the CPP peaks around the time of the response, there are likely differences with respect to both the latency and/or amplitude distributions across confidence ratings, which are unable to be cleanly dissociated in stimulus-locked ERPs (for further discussion see Ouyang et al., 2015; Feuerriegel et al., 2022).

Studies measuring pre-response CPP amplitudes (which avoid the issues described above) have yielded mixed results. Grogan et al. (2023) reported larger CPP amplitudes in trials with higher confidence ratings, both when choice and confidence were reported simultaneously and for delayed confidence judgments. However, Feuerriegel et al. (2022) reported that confidence was associated with amplitudes of a fronto-central component rather than the CPP (resembling components identified in Kelly & O’Connell, 2013; Burwell et al., 2019). This fronto-central effect also appeared to bias measures at parietal channels via volume conduction across the scalp (although this issue was accounted for in Grogan et al., 2023). It remains to be seen whether associations between response-locked CPP amplitudes and confidence reliably replicate across different decision-making tasks, or if there are a subset of conditions under which these correlations are observed (discussed in Grogan et al., 2023).

### 1.2. Postdecisional ERP Correlates of Confidence

Researchers have also reported associations between confidence ratings and amplitudes of the postdecisional Pe component. The Pe is typically measured around 200-400 ms after the time of the first-order decision at the same centro-parietal electrodes used to measure the CPP (e.g., Boldt & Yeung, 2015). Larger (i.e., more positive-going) Pe amplitudes are observed when participants are aware of making an error (Ridderinkhof et al., 2009; Steinhauser & Yeung, 2010; Wessel et al., 2011), and the morphology of this component closely resembles that of the CPP in the lead-up to error detection reports following a first-order decision (Murphy et al., 2015). Larger Pe amplitudes have also been observed in trials with lower confidence ratings (e.g., Boldt & Yeung, 2015; Desender et al., 2019a; Rausch et al., 2020). Based on these findings, Desender et al. (2021b) proposed a model in which the Pe component tracks the degree of postdecisional evidence accumulated *against* the first-order decision (i.e., the extent of evidence in favour of having made an error) that determines one’s degree of confidence. Importantly, their model specifies a monotonic, negative-going association between Pe amplitudes and confidence ratings spanning the range of “certainly wrong” to “unsure” to “certainly correct”.

Interpretation of these findings, however, is complicated by measurement issues relating to ERP baselines. Landmark findings of confìdence-Pe component associations have used pre-response baselines for their primary analyses (e.g., Boldt & Yeung, 2015; Desender et al., 2019b). These pre-response baseline windows overlap in time with observed differences in pre-response ERPs across confidence ratings, visible at the same centro-parietal electrodes (e.g., Feuerriegel et al., 2022; Grogan et al., 2023). Consequently, ERP differences already present during the pre-response baseline window would be artefactually propagated with opposite polarity to the post-response time window due to the baseline subtraction procedure (for examples see Feuerriegel & Bode, 2022). Although Boldt and Yeung (2015) reported similar results across pre-stimulus and pre-response baselines, others have reported that postdecisional ERP amplitude associations with confidence disappear when instead applying pre-stimulus baselines (Desender et al., 2019b; Feuerriegel et al., 2022; Grogan et al., 2023). Notably, Feuerriegel et al. (2022) reported that, when using a pre-stimulus baseline, Pe amplitudes were specifically associated with confidence ratings indicating certainty in having committed an error (across the range of “surely incorrect” to “unsure” ratings), but not certainty in having made a correct response (across the range of “unsure” to “surely correct”). Grogan et al. (2023) did observe postdecisional ERP waveforms that covaried with confidence ratings when using pre-stimulus baselines, however these were only observed in conditions where additional, decision-relevant stimuli were presented immediately after the time of the perceptual judgment in each trial. Rather than an evidence accumulation process based on residual representations of sensory evidence or iconic memory (as proposed in Murphy et al., 2015; Desender et al., 2021b), this may have instead reflected the additional accumulation of the sensory evidence provided by the postdecisional stimuli. In conditions where no stimuli were presented after the initial choice, associations between confidence and Pe amplitudes were not observed.

As the model of Desender and colleagues (2021b) provides an elegant, unified account of the Pe linking error detection, confidence, and changes of mind, further systematic investigation is necessary to determine the contexts in which the Pe does (and does not) vary with confidence in decision-making tasks.

### 1.3. Decorrelating Stimulus Discriminability and Confidence

There is another issue common to both the CPP- and Pe-related findings described above. In previous studies, higher average confidence ratings have been reported in conditions of higher stimulus discriminability (e.g., Rausch et al., 2020; Feuerriegel et al., 2022; Grogan et al., 2023), or in subsets of trials with higher accuracy (e.g., Boldt & Yeung, 2015). It has been proposed that, in such cases, neural correlates of confidence may (at least partly) reflect co-occurring differences in stimulus discriminability or task difficulty rather than one’s degree of confidence *per se* (discussed in Lau & Passingham, 2006; Odegaard et al., 2018; Dou et al., 2023).

However, confidence and accuracy can be (at least partly) dissociated. Recent studies using two-choice discrimination tasks have manipulated task difficulty (e.g., the difference in brightness between two squares in a comparative brightness judgment task, here termed relative evidence) as well as the overall intensity of a relevant sensory attribute (the overall brightness of the two squares, here termed absolute evidence). According to Weber’s law, increasing the level of absolute evidence (while keeping relative evidence constant) reduces stimulus discriminability (Geisler, 1989). However, increases in absolute evidence also lead to faster responses and higher levels of reported confidence despite worse accuracy in these conditions (e.g., Zylberberg et al., 2012; Koizumi et al., 2015; Peters et al., 2017; Odegaard et al., 2018; Samaha & Denison, 2022; Ko et al., 2022). Manipulations of stimulus luminance have also produced similar effects on confidence and accuracy in recognition memory tasks (Busey et al., 2000). This implies that absolute evidence influences both decision-making and metacognition, which presents an opportunity to probe neural correlates of confidence that are not strongly correlated with accuracy.

Recently, Dou et al. (2023) manipulated both relative and absolute evidence and reported consistent associations between CPP build-up rates (i.e., rising slopes) and confidence across three experiments. This was found even when statistically controlling for levels of relative evidence, suggesting that pre-decisional neural correlates of confidence may be distinct from ERP components that covary with task difficulty. However, they did not analyse CPP pre-response *amplitudes,* which are thought to index the *extent* of evidence accumulated, as opposed to the CPP build-up rate which is linked to the *rate* of evidence accumulation (O’Connell et al., 2012; Kelly & O’Connell, 2013). The former, which is directly relevant to racing accumulator models, remains to be tested.

### 1.4. Testing for Pre- and Postdecisional Correlates of Confidence

Despite extensive efforts to characterise the ERP correlates of confidence, a large body of findings are complicated by ERP measurement issues, and recent findings have not been consistently replicated across studies. To further test the generalisability of pre- and postdecisional ERP correlates of confidence across decision contexts, we adapted the brightness judgment design of Ko et al. (2022) and recorded EEG. In this task, participants judged which of two luminance-varying squares was brighter on average and provided confidence ratings after a brief interval. We manipulated the average luminance difference (termed relative evidence in Ko et al., 2022) and the overall luminance of both squares (absolute evidence). As observed in previous work (e.g., Ko et al., 2022; Dou et al., 2023) higher levels of absolute evidence led to worse accuracy but higher confidence ratings. This allowed us to assess neural correlates of confidence ratings that were partially decorrelated from relative evidence and task performance (as done in Dou et al., 2023). Here, we tested for associations between confidence ratings and pre-decisional CPP amplitudes (linked to the extent of evidence accumulation at the time of a decision) as well as postdecisional Pe amplitudes (linked to the extent of evidence accumulated in favour of detecting an error).

## 2. Method

### 2.1. Participants

We recruited 36 university students with normal or corrected-to-normal vision. We report how we determined our sample size, all data exclusions (if any), all inclusion/exclusion criteria, whether inclusion/exclusion criteria were established prior to data analysis, all manipulations, and all measures in the study. Our target sample size (prior to exclusions) was comparable to Feuerriegel et al. (2022, n=35) who reported ERP correlates of confidence. We did not have a specific target effect size, however we note that our sample size is larger than those of many previous studies (e.g., n=l6 in Boldt & Yeung, 2015; n=25 in Rausch et al., 2020). We excluded two participants who failed to report confidence ratings in more than 20% of all trials, three for overall accuracy lower than 55%, and one for reporting the same confidence level in more than 90% of trials whereby confidence was reported (same exclusion criteria as in Ko et al., 2022). Two further participants were excluded due to frequent, long-duration blinks and eye movements during the experimental trials as identified by visual inspection of the data. This criterion was not established prior to data analysis but was judged to be a strong indicator that those participants were not performing the task in the intended manner. The final sample included 28 participants (aged 18-39, M=26, SD=6,16 female). This sample size was comparable to recent studies that reported associations between CPP and Pe amplitudes and confidence (e.g., Feuerriegel et al., 2022; Grogan et al., 2023). Participants were reimbursed 30 AUD for their time. This study was approved by the Human Research Ethics Committee of the Melbourne School of Psychological Sciences (ID 1954641.2).

### 2.2. Stimuli and Task

Participants completed a comparative brightness judgment task in which two flickering greyscale squares were concurrently presented in each trial. Both squares were 70 × 70 pixels in size and were positioned at equal distances from the centre of the screen, separated from each other by 180 pixels. Stimuli were displayed against a black background on a Sony Trinitron Multiscan G42O CRT Monitor (1280 x 1024 pixels; 75 Hz refresh rate) gamma-corrected using a ColorCAL MI<II Colorimeter (Cambridge Research Systems). Stimuli were presented using Psychtoolbox-3 (Brainard, 1997; Kleiner et al., 2007) running in MATLAB 2018b (The Mathworks). Code used for stimulus presentation will be available at osf.io/7xv3m at the time of publication.

Within each trial (depicted in Figure 1A), the luminance of each flickering square changed with each monitor refresh (every 13.3 ms). Luminance values of each square were randomly sampled from a pair of truncated normal distributions which differed in average luminance. The brighter square was on the left in half of trials and on the right in the other half of trials.

**Figure 1.**
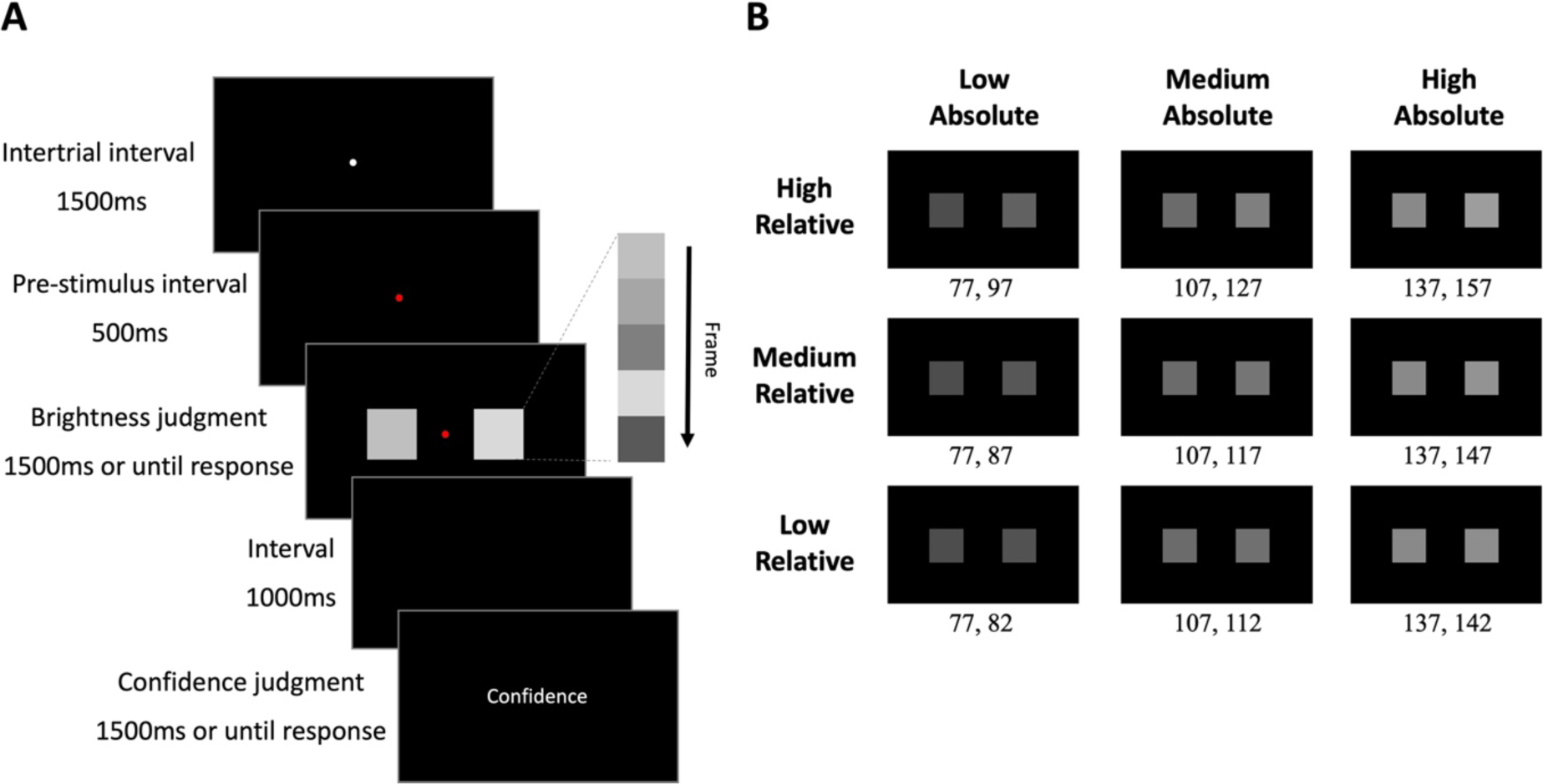
Trial diagram and stimuli. (A) In each trial, two flickering squares of differing average luminance were presented. Each square changed in luminance with each monitor refresh (every 13.3 ms). Participants indicated the square that appeared brighter on average. Following this judgment, participants reported their decision confidence using a 7-point scale while the word “confidence” was presented on the screen. (B) Average luminance values (in RGB) for each square in each condition. Luminance values differed across conditions with respect to the difference between squares (low/medium/high relative evidence) and the overall luminance across both squares (low/medium/high absolute evidence).

Mean RGB values for the overall brighter and darker squares differed across conditions (values specified in Figure 1B, same as in Ko et al., 2022). The truncated normal distributions had standard deviations of 25.5 RGB values and were truncated at ± 1 SD from their means as in Ratcliff et al. (2018) and Ko et al. (2022).

Each condition varied with respect to the difference in average luminance across the brighter and darker squares (here termed the degree of *relative evidence)* and the overall luminance of both squares (termed *absolute evidence,* using terminology in Teodorescu et al., 2016; Turner et al., 2021b; Ko et al., 2022). There were nine conditions in total, including all combinations of the three levels of relative and absolute evidence levels. These combinations of relative and absolute evidence produced a range of confidence ratings in Ko et al. (2022) and were used to elicit variation in confidence ratings here.

### 2.3. Procedure

After giving written consent and receiving task instructions, participants were seated in a dark testing booth 70 cm from the computer monitor. They completed a training session while the experimenter stayed in the testing booth to ensure task comprehension. Participants then completed the main experiment alone. No part of the study procedures was pre-registered prior to the research being conducted.

In each trial (depicted in Figure 1A) a white fixation dot was presented for 1,500 ms. The dot changed to red for 500 ms to signal the imminent appearance of the flickering squares. Following this, the squares appeared. Participants indicated which square appeared brighter on average by pressing one of two buttons on a seven-button Cedrus response pad (RB-740, Cedrus Corporation) using their left and right index fingers. Stimuli were presented for a maximum of 1,500 ms and disappeared immediately after a response was made. Participants were instructed to respond as quickly as possible.

Participants were then presented with a blank screen for 1,000 ms and subsequently rated their degree of confidence using the same seven-button response pad. During this period, the word ‘Confidence’ was presented in the centre of the screen. Participants indicated their confidence on a seven-point scale with options “surely incorrect” (1), “probably incorrect” (2), “maybe incorrect” (3), “unsure” (4), “maybe correct” (5), “probably correct” (6) and “surely correct” (7). The midpoint rating of “unsure” signified that they were unsure whether the brightness judgment was correct or incorrect (i.e., they indicated that they were guessing). They were also instructed to make their confidence rating as quickly as possible within 1,500 ms from confidence rating stimulus onset. No confidence rating was required if the brightness judgment was “too slow” (>1,500 ms RT) or “too quick” (<250 ms RT). In this case, only the respective “too early/slow” feedback was presented for 1,500 ms, and then the next trial began.

The experiment included 1,008 experimental trials equally allocated across 14 blocks of 72 trials each. An equal number of trials from all conditions were randomly interleaved within each block. Each block was followed by a self-terminated rest period.

Before the experiment, participants completed a training block of 36 trials without making confidence ratings. Instead, they received feedback on their accuracy after each trial to familiarize themselves with the brightness judgment task. Participants then completed another training block of 36 trials in which both brightness and confidence judgments were required. No performance feedback was given in this training, and a confidence rating scale displaying each option was presented on the screen for 1,500 ms. Participants were instructed that during the main experiment, the visual presentation of the scale would be removed, and the confidence judgment would only be prompted by the word “Confidence”.

### 2.4. Task Performance Analyses

To characterise patterns of task performance we first examined effects of relative and absolute evidence manipulations on brightness judgment accuracy, RTs and confidence ratings. Analyses of RTs and confidence ratings were done separately for trials with correct and erroneous brightness judgments (as done by Fleming et al., 2018; Ko et al., 2022). Code used for analyses will be available at osf.io/7xv3m at the time of publication. No part of the study analyses was pre-registered prior to the research being conducted.

We fit generalised linear mixed effects models using the R package Ime4 (Bates et al. 2015). We included relative evidence, absolute evidence, and their interaction as fixed effects and participant as a random intercept in all models. We fit models specifying binomial distributions with a logit function to model accuracy, gamma distributions with an identity function for RTs (as recommended by Lo & Andrews, 2015), and gaussian distributions with an identity function for confidence (as done by Fleming et al., 2018). Model equations and outputs are presented in the Supplementary Material.

We additionally performed frequentist and Bayesian post-hoc paired-samples t tests using JASP vθ.17.1 (JASP Core Team, Cauchy prior distribution, width 0.707, default settings) to determine whether average confidence ratings were higher for trials with correct as compared to erroneous responses. We performed this analysis (using pooled data across relative and absolute evidence conditions) for all trials, and for the subset of trials that contributed to averaged ERPs for correct responses and errors.

### 2.5. EEG Data Acquisition and Processing

EEG data were recorded using a Biosemi Active Two system with 64 active electrodes at a sampling rate of 512 Hz. Recordings were grounded using common mode sense and driven right leg electrodes (http://www.biosemi.com/faq/cms&drl.htm). Six external electrodes were additionally included: two behind the left and right mastoids, two placed 1 cm from the outer canthi, and one above and one below the right eye.

EEG data were processed using EEGLab v.2022.0 (Delorme & Makeig, 2004) in MATLAB (Mathworks). First, we identified excessively noisy channels by visual inspection (median number of bad channels = 0, range 0-9) and excluded these from average reference calculations and Independent Components Analysis (ICA). Sections with large artefacts were also manually identified and removed.

We then re-referenced the data to the average of all channels, low-pass filtered the data at 30 Hz (EEGLab Basic Finite Impulse Response Filter New, default settings), and removed one extra channel (AFz) to correct for the rank deficiency caused by the average reference. We processed a copy of this dataset in the same way and additionally applied a 0.1 Hz high-pass filter (EEGLab Basic FIR Filter New, default settings) to improve stationarity for the ICA. We then ran ICA on the duplicate dataset using the RunICA extended algorithm (Jung et al., 2000). Independent component information was copied to the unfiltered dataset (e.g., as done by Feuerriegel et al., 2018). Components associated with muscular and ocular activity were identified and removed based on the guidelines in Chaumon et al. (2015). Previously removed channels, including AFz, were then interpolated from the cleaned dataset using spherical spline interpolation.

The resulting EEG data were then segmented from −100 to 2,200 ms relative to flickering square onset. Segments were baseline corrected using the pre-stimulus interval. Epochs with amplitudes exceeding ±150 µV at any scalp electrode were excluded from further analyses. Across participants, the number of epochs included in EEG analyses ranged from 335 to 982 (median = 744). From this data, we derived response-locked segments ranging from −500 to 700 ms relative to the brightness judgment button press.

Two versions of the response-locked data were created. The first used the same pre-stimulus baseline as the stimulus-locked segments. The second used a pre-response baseline spanning −100 to 0 ms relative to the button press response. This was done to investigate how the use of pre-response baselines (commonly used in this area of research) influences measures of post-decisional ERP components such as the Pe (see Feuerriegel et al., 2022, Feuerriegel & Bode, 2022).

### 2.6. ERP Component Amplitude Analyses

#### 2.6.1. CPP amplitude analyses

We measured response-locked CPP amplitudes as the average amplitude between −130 and −70 ms relative to the response, averaged across parietal electrodes Pz, Pl, P2, CPz, and POz (same time window as Steinemann et al., 2018; Feuerriegel et al., 2021, 2022). For these analyses, we used a pre-stimulus baseline. To link our results to previous work using stimulus-locked ERPs (e.g., Gherman & Philiastides, 2018; Rausch et al., 2020), we also measured stimulus-locked CPP amplitudes as the average amplitude within the time window of 350-500 ms from flickering square onset.

For CPP amplitude analyses we compared correct and erroneous responses using paired-samples frequentist and Bayesian t-tests as implemented in JASP vθ.17.1 (JASP Core Team, Cauchy prior distribution, width 0.707, default settings). We additionally fitted linear regression models using MATLAB to predict CPP mean amplitudes based on confidence ratings within the range spanning “maybe correct” (5) to “surely correct” (7) for each participant separately, as done by Feuerriegel et al. (2022). As the “guessing” (4) rating may also be considered as the lowest certainty condition, we also included complementary analyses that included this rating. For these analyses, we included both trials with correct responses and errors. The resulting Beta coefficients (regression model slopes) were tested at the group level using one-sample frequentist and Bayesian t-tests (as done by Feuerriegel et al., 2022). We did not perform analyses using confidence ratings between “surely incorrect” and “maybe incorrect” (i.e., ratings indicating an error had occurred) because there were insufficient numbers of trials for such restricted analyses (median number of trials per participant = 33.5, and 9 participants with < 20 trials). Please note that, although there were substantial numbers of trials with errors, in those trials, participants often provided confidence ratings indicating that they had made a correct response (as also observed in Ko et al., 2022).

To test for associations between CPP amplitudes and confidence while controlling for relative evidence, we performed confìdence-CPP amplitude regression analyses using the methods described above for trials within each relative evidence condition separately (similar to analyses in Tagliabue et al., 2019). We used restricted ranges of confidence ratings spanning “maybe correct” to “surely correct”. In a complementary analysis we averaged each of these beta (regression slope) estimates across participants to derive overall measures of associations with confidence.

We also assessed effects of relative and absolute evidence independently of effects of confidence, to determine whether CPP amplitudes also covaried with the quality of information provided by the stimulus (similar to Odegaard et al., 2018; Tagliabue et al., 2019). We used the same regression analysis methods to test for associations with relative evidence level (low, medium, high) when holding confidence constant, for “maybe correct”, “probably correct” and “surely correct” ratings separately. Beta values were also averaged across confidence rating conditions to derive more general estimates of relative evidence effects. We repeated these analyses using absolute evidence level (low, medium, high) as a predictor of CPP amplitudes while holding confidence constant.

#### 2.6.2. Pe amplitude analyses

For Pe amplitude measures we baseline-corrected single-trial ERPs using pre-stimulus and pre-response baselines in separate analyses. We measured Pe amplitudes as the mean amplitude within 200-350 ms following the response, using the same set of parietal electrodes as for the CPP (same time window as Nieuwenhuis et al., 2001; Di Gregorio et al., 2018; Feuerriegel et al., 2022). Paired-samples t-tests were conducted to compare Pe amplitudes across trials with correct and erroneous responses. Within-subject regressions were performed using the predictor of confidence as described above. We did not find associations between Pe amplitudes and confidence in the data using pre-stimulus baselines, and so we did not conduct additional analyses controlling for relative or absolute evidence.

## 3. Results

### 3.1. Task Performance

Both relative and absolute evidence manipulations produced intended effects on accuracy, RTs, and confidence ratings, consistent with Ko et al. (2022, see also Ratcliff et al., 2018; Turner et al., 2021b). Task performance for each relative and absolute evidence level combination is shown in Figure 2. Accuracy was higher in conditions with higher relative evidence (i.e., larger mean luminance differences between the squares, p < .001) and lower in conditions of higher absolute evidence (i.e., higher overall luminance across the two squares, p < .001, see Figure 2A). RTs for correct and error trials were faster in conditions of higher absolute evidence and higher relative evidence (p’s < .01 for all main effects, Figure 2B-C) except for RTs in error trials where relative evidence did not have a significant effect. Higher confidence ratings were made in conditions of higher relative evidence and higher absolute evidence for trials with correct responses (p’s < .001, Figure 2D). This was despite lower accuracy in higher absolute evidence conditions and consistent with findings in Ko et al. (2022). In trials with errors, confidence ratings were higher in conditions of higher absolute evidence and lower relative evidence (p’s < .001, Figure 2E). Model outputs and the full set of statistical results (including interaction effects) are reported in the Supplementary Material. Associations between confidence, accuracy, and RTs are plotted in Supplementary Figure S1.

**Figure 2.**
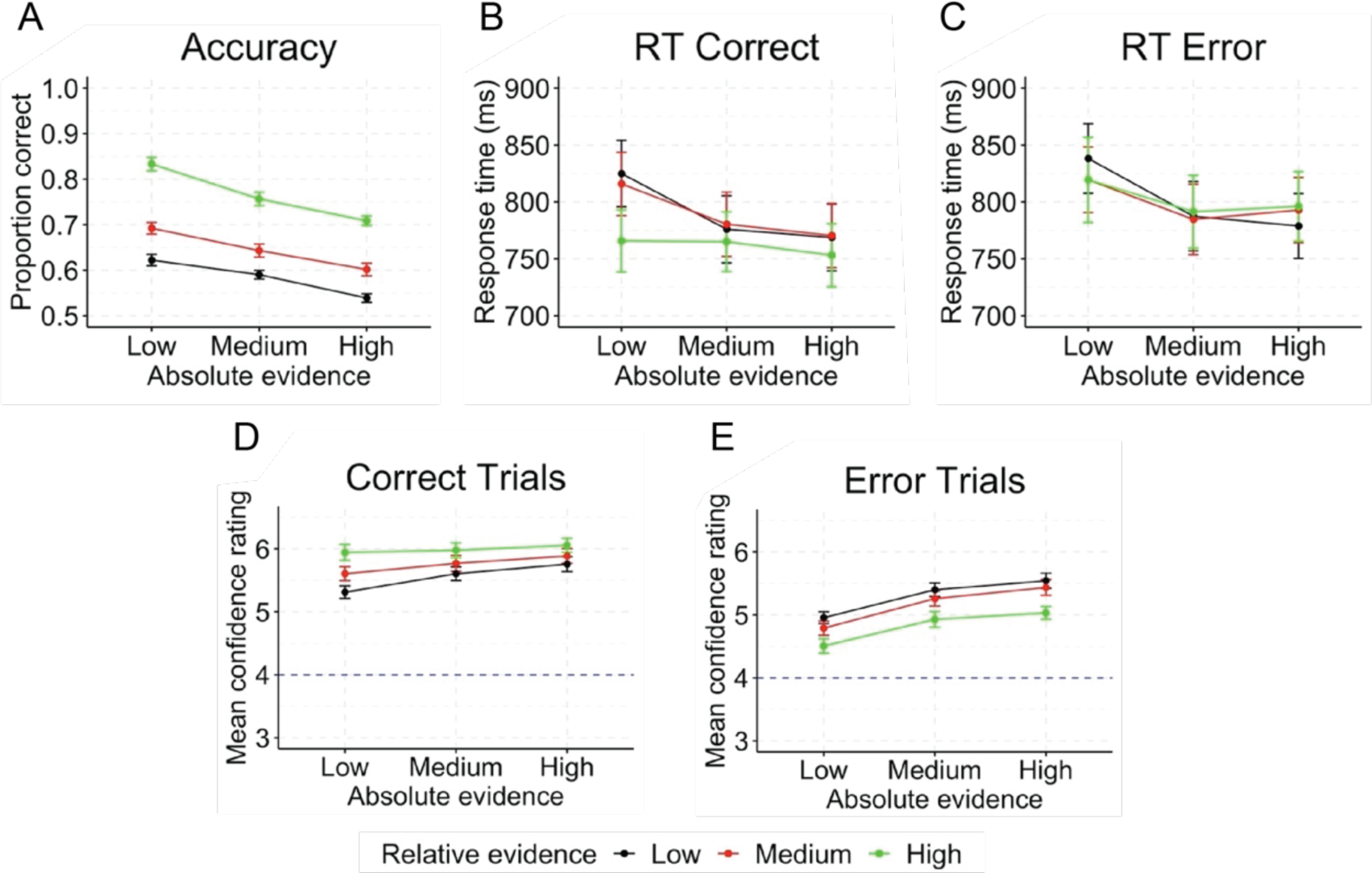
Accuracy, mean RTs, and mean confidence ratings for different combinations of relative and absolute evidence levels. (A) Decision accuracy (proportion correct). (B) Mean RTs for correct trials. (C) Mean RTs for error trials. (D) Mean confidence ratings for correct trials. (E) Mean confidence ratings for error trials. Confidence ratings were measured on a scale ranging from i (surely incorrect) to 7 (surely correct) with a mid-point of 4 indicating guessing. The dotted line indicates the mid-point of the scale. Error bars represent standard errors.

We also performed post-hoc paired-samples t tests to assess whether confidence ratings differed across trials with correct and erroneous responses. When including all trials, confidence was higher on average in trials with correct responses, t(27) = 12.09, p < .001, BF_10_ = 3.88 * 10^9^. However, when only including the subset of trials that contributed to averaged ERPs for correct and error responses, we did not observe differences in average confidence ratings, t(27) = 0.90, p = .374, BF_10_ = 0.29.

### 3.2. CPP Amplitudes During Decision Formation

First, we analysed CPP amplitudes time-locked to stimulus onset as done in some previous studies (e.g., Rausch et al., 2020). We did not find significant differences in stimulus-locked CPP amplitudes across trials with correct responses and errors, t(27) = 0.13, p = .898, BF_10_ = 0.20 (Figure 3A). We also did not find evidence for associations with confidence across the range of “maybe correct” to “surely correct” ratings, t(27) = −0.43, p = .673, BF_10_ = 0.22 (Figure 3C). Results of complementary analyses that included the “guessing” trials as the lowest confidence rating did not yield associations with CPP amplitudes, t(27) = −0.27, p = .787, BF_10_ = 0.21.

**Figure 3.**
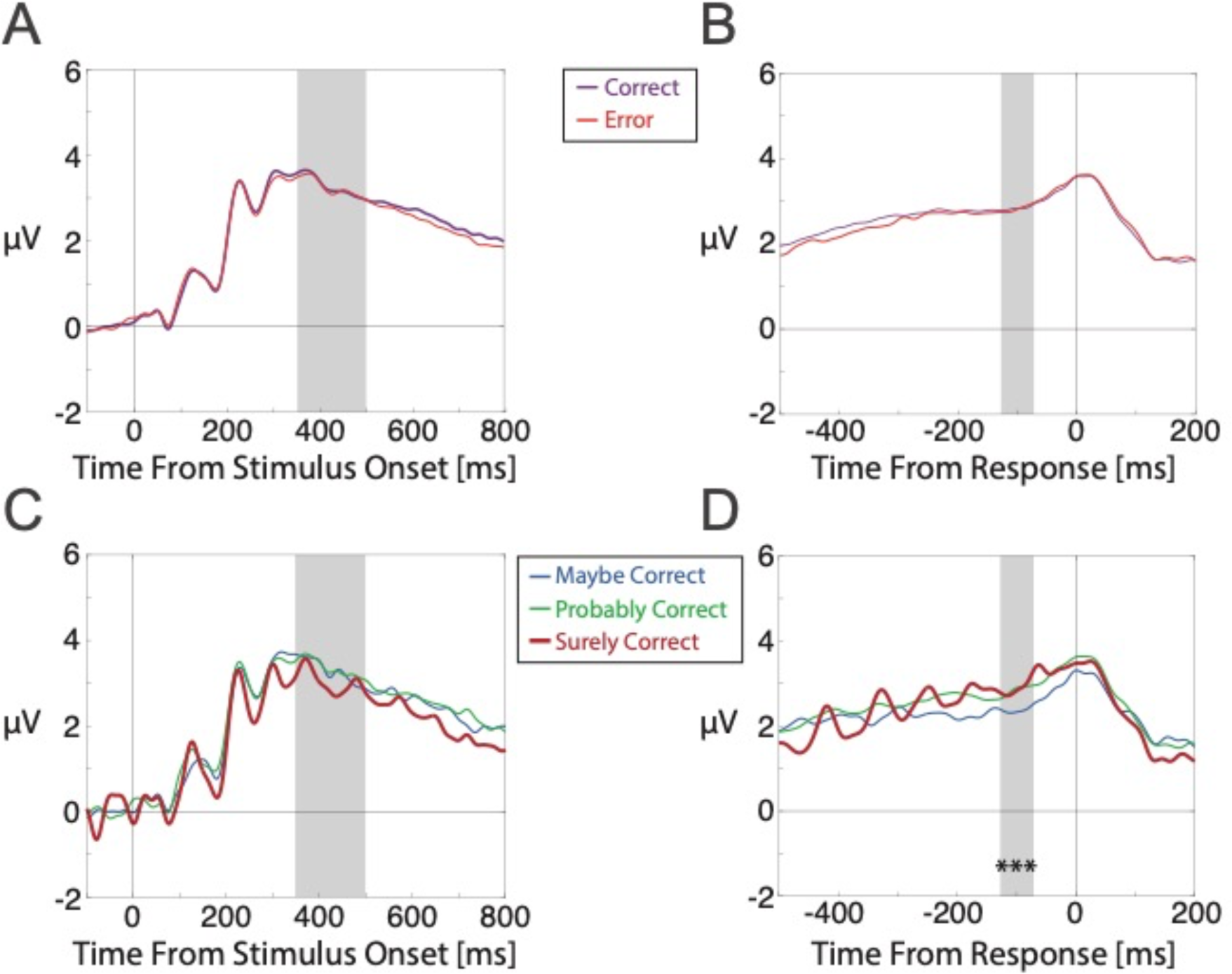
Stimulus- and response-locked ERPs averaged over parietal channels Pz, Pl, P2, CPz, and POz. Stimulus-locked ERPs are plotted in the left column. Response-locked ERPs are plotted in the right column. A, B) ERPs for correct responses and errors. B, D) ERPs for confidence ratings ranging from “maybe correct” to “surely correct”. Grey shaded regions denote the mean amplitude measurement time windows for the CPP. Asterisks denote statistically significant associations between confidence ratings and ERP component amplitudes (*** denotes p < .001 for the regression analysis using confidence ratings ranging between “maybe correct” to “surely correct”).

We then repeated these analyses for CPP amplitudes time-locked to the response. We did not find differences in pre-response CPP amplitudes across correct responses and errors, t(27) = 0.27, p = .790, BF_10_ = 0.21 (Figure 3B). Here, please note that average confidence ratings did not significantly differ between correct responses and errors when analysing trials that contributed to these averaged ERPs, as reported above. To assess associations between pre-response CPP amplitudes and certainty in having made a correct response as reported in Feuerriegel et al. (2022) and Grogan et al. (2023), we performed regression analyses using restricted ranges of confidence ratings, spanning “maybe correct” (5) to “surely correct” (7). Here, omitting the “guessing” (4) trials further removed any ambiguity about whether this option could have been chosen in the case of a lapse of attention. CPP amplitudes were positively associated with confidence, t(27) = 4.20, p < .001, BF_10_ = 111.81 (Figure 3D). Scalp maps of group-averaged beta values (i.e., regression model slopes) indicated that associations between CPP amplitudes and confidence were most prominent over parietal channels (Supplementary Figure S2A). We also ran complementary analyses that included the “guessing” trials as the lowest certainty rating. CPP amplitudes remained associated with confidence in these analyses, t(27) = 2.25, p = .032, BF_10_ = 1.74.

#### 3.2.1. Associations with confidence when controlling for relative evidence

We also tested for associations between pre-response CPP amplitudes and certainty in having made a correct response while controlling for effects of relative evidence. To do this, we fit regression models separately for trials within each relative evidence level. We again restricted our analyses to confidence judgments ranging between “maybe correct” (5) and “surely correct” (7), given the low trial numbers for each analysis in the lower confidence range after splitting by relative evidence level. CPP amplitudes were larger in trials with higher confidence ratings compared to lower confidence ratings, for conditions of low relative evidence, t(27) = 2.42, p = .023, BF_10_ = 2.33, medium relative evidence, t(27) = 4.02, p < .001, BF_10_ = 72.62, and high relative evidence, t(27) = 3.71, p < .001, BF_10_ = 35.18. Associations with confidence were also found when averaging beta estimates across low, medium, and high relative evidence for each participant, t(27) = 3.87, p < .001, BF_10_ = 50.75.

#### 3.2.2. Effects of relative and absolute evidence when controlling for confidence

We also tested for associations between the level of relative evidence and pre-response CPP amplitudes while keeping confidence ratings constant. This was done for trials with “maybe correct”, “probably correct” and “surely correct” ratings in separate analyses. We did not find associations between CPP amplitudes and relative evidence level for “maybe correct”, t(27) = −0.73, p = .473, BF_10_ = 0.26, “probably correct”, t(27) = 0.03, p = .975, BF_10_ = 0.20, or “surely correct” ratings, t(27) = 0.31, p = .760, BF_10_ = 0.21, nor when averaging beta values across each of these analyses within each participant, t(27) = −0.36, p = .723, BF_10_ = 0.21.

We performed the same analyses using absolute evidence as a predictor. We did not find associations between CPP amplitudes and absolute evidence level for “maybe correct”, t(27) = 1.23, p = .229, BF_10_ = 0.40, “probably correct”, t(27) = −0.04, p = .968, BF_10_ = 0.20, or “surely correct” ratings, t(27) = 0.10, p = .925, BF_10_ = 0.20, nor when averaging beta values across each of these analyses within each participant, t(27) = 0.86, p = .397, BF_10_ = 0.28.

Taken together, these results indicate that CPP amplitudes covaried with confidence rather than levels of relative or absolute evidence.

### 3.3. Post-Decisional Pe Component Amplitudes

We analysed Pe component amplitudes using both pre-stimulus and pre-response ERP baselines in separate analyses, as done in Feuerriegel et al. (2022) and Grogan et al. (2023). This was done to systematically assess whether the use of pre-response baselines can artificially produce associations between Pe amplitudes and confidence, in cases where there are already ERP differences during the pre-response baseline window (e.g., Feuerriegel & Bode, 2022).

#### 3.3.1. Analyses using pre-stimulus baselines

When using pre-stimulus baselines (which circumvent issues with propagating potential pre-response ERP differences into the Pe window), we did not observe Pe amplitude differences between errors and correct responses, t(27) = −0.24, p = .81, BF_10_ = 0.21 (Figure 4A). Pe amplitudes were not associated with decision confidence, t(27) = 0.417, p = .680, BF_10_ = 0.22 (Figure 4C). We also ran complementary analyses that included the “guessing” trials as the lowest confidence rating. Pe amplitudes were not associated with confidence in these analyses, t(27) = −1.02, p = .316, BF_10_ = 0.32. Because there was no indication of Pe amplitudes covarying with confidence, we do not report additional analyses controlling for relative or absolute evidence here.

**Figure 4.**
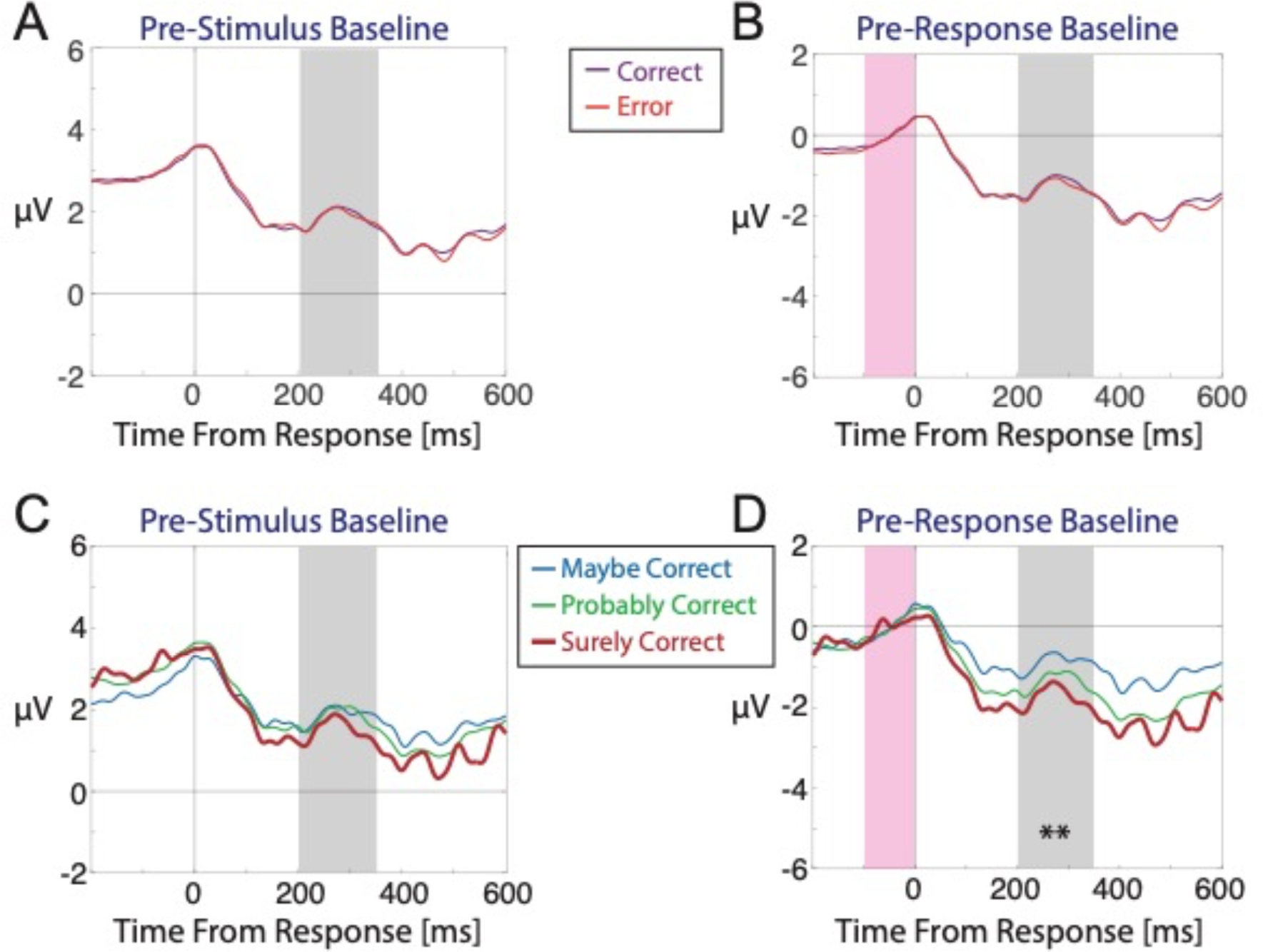
Group-averaged ERPs following keypress responses at electrodes Pz, Pl, P2, CPz, and POz, corrected using either a pre-stimulus baseline (left column) or a pre-response baseline (right column). A, B) Trials with correct responses and errors. C, D) ERPs for confidence ratings ranging from “maybe correct” to “surely correct”. In all plots the grey shaded area denotes the 200-350 ms time window used to measure the Pe component. The shaded magenta area denotes the pre-response baseline time window. Asterisks denote statistically significant associations between confidence ratings and ERP component amplitudes (** denotes p < .01 for the regression analysis using confidence ratings from “maybe correct” to “surely correct” included).

#### 3.3.2. Analyses using pre-response baselines

When using pre-response baselines (which, however, might propagate potential pre-existing ERP differences into the Pe window), we did not identify differences in Pe amplitudes following errors compared to correct responses, t(27) = −1.13, p = .268, BF_10_ = 0.36 (Figure 4B). However, there were clear associations between Pe amplitudes and confidence ratings, t(27) = −2.87, p = .008, BF_10_ = 5.61 (Figure 4D), such that more positive-going amplitudes were observed in trials with lower confidence ratings. We also ran complementary analyses that included the “guessing” trials as the lowest confidence rating. Pe amplitudes remained associated with confidence in these analyses, t(27) = −3.71, p < .001, BF_10_ = 35.72. However, as explained above, these statistically significant associations are most likely due to differences already present before the response (during the CPP measurement window, Figure 3D) that largely overlap with the pre-response baseline window.

In summary, these results show that in our experiment, in which error detection was rare, the Pe did not reliably reflect differences between errors and correct responses. When avoiding confounds relating to pre-response ERP differences, the Pe also did not reflect differences in confidence judgments.

## 4. Discussion

To test for electrophysiological correlates of confidence we presented participants with a comparative brightness judgment task. We elicited a wide range of confidence ratings by varying the difference in average luminance across the two flickering squares (relative evidence), and the overall brightness of both squares (absolute evidence). CPP amplitudes during decision formation were positively associated with confidence, and specifically certainty in having made a correct response. Importantly, this was not simply a consequence of covariation between relative evidence (task difficulty) and confidence ratings. Our findings complement reports of parietal (Grogan et al., 2023) and fronto-central (Feuerriegel et al., 2022) correlates of confidence during decision formation, suggesting that critical computations that inform confidence judgments occur during this time window. However, we did not find associations between confidence and Pe component amplitudes following each decision, except when these were likely to be artificially produced using pre-response baselines (as in Feuerriegel et al., 2022; Grogan et al., 2023). We show that, in some decision contexts, postdecisional ERP trajectories do not necessarily covary with confidence. Instead, we propose that postdecisional ERP dynamics depend on whether postdecisional evidence accumulation is relied upon to inform confidence judgments, and also specific conditions such as the reporting of decision errors (e.g., Murphy et al., 2015; Feuerriegel et al., 2022) or continued integration of additional sensory evidence after a decision is made (Grogan et al., 2023).

### 4.1. ERP Correlates of Confidence During Decision Formation

We found larger (more positive) CPP amplitudes during decision formation when participants made higher confidence ratings (similar to Feuerriegel et al., 2022; Grogan et al., 2023). This association persisted even when controlling for relative evidence, which often covaries with confidence across a variety of decision-making tasks (e.g., Philiastides et al., 2014; Fleming et al., 2018; Feuerriegel et al., 2022; Grogan et al., 2023). The absolute evidence manipulation in our paradigm allowed us to elicit a range of confidence judgments at each level of relative evidence. We found that CPP amplitudes were specifically associated with confidence rather than relative evidence in our data (similar to electrocorticography results in Peters et al., 2017) as also found in relation to perceptual awareness ratings (Tagliabue et al., 2019). Confìdence-ERP amplitude correlations were strongest over parietal channels rather than the more fronto-central distribution identified in Feuerriegel et al. (2022), suggesting that this effect does reflect variation in the CPP component.

Based on this interpretation, it is plausible that these effects reflect modulations of the CPP component, and by extension differences in the degree of (unsigned) evidence accumulated at the time of the decision (consistent with Grogan et al., 2023). Although a defining feature of CPP component morphology is that it typically reaches a fixed amplitude prior to the response, as consistent with standard implementations of the Diffusion Decision Model (O’Connell et al., 2012; Kelly & O’Connell, 2013), situations have been identified in which pre-response amplitudes systematically vary across conditions (e.g., Steinemann et al., 2018; Kelly et al., 2021; Feuerriegel et al., 2021). Assuming that the CPP does trace the degree of (unsigned) evidence in favour of the chosen option in each trial, modulations of pre-response amplitudes would be consistent with racing accumulator models of confidence that specify more accumulated evidence in favour of the chosen option in trials with higher confidence ratings (e.g., Vickers, 1979; Vickers & Packer, 1982; see also Smith et al., 2022). Although processes occurring during decision formation are unlikely to be the sole source of information used to guide confidence judgments (e.g., Fleming & Daw, 2017; Turner et al., 2021a; Desender et al., 2021a), this interpretation would suggest that evidence accumulation dynamics, which can be traced using neural measures, are important to consider when modelling confidence judgments.

This interpretation also has implications for post-decisional evidence accumulation models of confidence (e.g., Pleskac and Busemeyer, 2010; Moran et al., 2015; Desender et al., 2021b), most of which specify a fixed decision boundary for the initial choice to preserve model identifiability and prevent over-fitting (for a more flexible model see van den Berg et al., 2016b). If the extent of evidence accumulation at the time of the first-order choice varies across confidence judgments (as specified in racing accumulator models), then the fixed decision boundary assumption of many postdecisional locus models is false. If this is the case, it is likely that (at least part of) the variance captured by postdecisional processes in these models is due to processes that occur during the first-order judgment (e.g., Turner et al., 2022). To account for this, hybrid decisional/postdecisional process models could be developed to better delineate the consequences of computations occurring over each time window. However, model identifiability should be ascertained (e.g., using parameter recovery methods, Miletić et al., 2017; Evans et al., 2020) before they are fitted to choice and confidence data.

Here, we note that specific features of our trial design may have enabled detection of pre-decisional correlates of confidence. Our findings are consistent with those in Grogan et al. (2023) when a blank screen was presented between choice and delayed confidence ratings, but not when postdecisional stimuli were presented that provided additional information about the accuracy of the first-order decision. As noted by Grogan and colleagues, presentation of additional, decision-relevant stimuli (or substantial postdecisional evidence accumulation, e.g., Resulaj et al., 2009) may weaken the relationship between processes during decision formation and subsequent confidence ratings that are provided after some delay.

We also note that the topographic distributions beta values in Supplementary Figure S2A (displaying associations between ERP amplitudes and confidence) are most prominent across a broad range of parietal channels, which does not quite match the midline parietal distribution of the CPP (visible in Supplementary Figures S2C). This suggests that, contrary to the clear midline parietal effect loci in Grogan et al. (2023), additional, overlapping ERP components may have contributed to the observed effects. Notably, Feuerriegel et al. (2022) observed a distinct, fronto-central component that influenced amplitudes at parietal channels via volume conduction (see also Kelly & O’Connell, 2013; Dmochowski & Norcia, 2015; Burwell et al., 2019 for a similar component), however an effect with such a frontal locus was not observed in our data. The close proximity of any additional effects to midline parietal channels makes it difficult to cleanly disentangle those from CPP component changes using standard current source density (CSD) estimation methods (e.g., Kayser and Tenke, 2006), and so we have not used CSD-transformed ERPs here. Future work should pay close attention to the topographies of effects and their correspondence to well-established spatial distributions of the CPP component (e.g., Kelly & O’Connell, 2013).

### 4.2. Postdecisional ERP Correlates of Confidence

In our experiment, Pe amplitudes did not covary with confidence ratings. However, the application of a pre-response baseline produced spurious associations with confidence (discussed in Feuerriegel et al., 2022; Feuerriegel & Bode, 2022). These findings are congruent with previous work that included analyses using both pre-stimulus and pre-response baselines (Desender et al., 2019b; Feuerriegel et al., 2022; Grogan et al., 2023; but see Boldt & Yeung, 2015). Although Feuerriegel et al. (2022) reported more positive-going Pe amplitudes for error trials with lower confidence ratings, this was specifically linked to certainty in having made an error. In other words, an association was found specifically when analysing the range of confidence ratings indicating that an error had occurred, similar to the “surely incorrect” to “maybe incorrect” range here. We did not obtain sufficient numbers of trials with error-indicating confidence ratings to evaluate the same effect in our dataset.

Our findings are relevant to the recently-proposed model of Desender et al. (2021b) that posits that postdecisional parietal ERP components (specifically the Pe) reflect a continued evidence accumulation process that determines confidence judgments. According to this model, two-alternative discrete choice decisions are initially made according to a double-bounded evidence accumulation process. After the first-order judgment is made, the ensuing metacognitive confidence judgment is proposed to reflect the degree of evidence accumulated in favour of having made an error. Importantly, the model of Desender et al. (2021b) predicts that the extent of accumulated postdecisional evidence will be reflected in the amplitude of the Pe component, and that the amplitude of the Pe component will show a monotonic, inverse relationship with decision confidence ratings spanning the range of “surely incorrect” to “surely correct” (as depicted in their Fig. 1C).

Contrary to these predictions, we did not observe such differences in Pe amplitudes across confidence ratings. Our findings indicate that, in some circumstances, confidence ratings are more closely associated with neural activity at the time of the decision rather than postdecisional ERPs. Notably, in our task any postdecisional evidence accumulation may not have strongly influenced confidence judgments because the stimuli were difficult to discriminate and fluctuated over time. This contrasts with Flanker or Stroop tasks in which the stimuli are highly visible, and participants can more easily detect their errors (e.g., Murphy et al., 2015). Postdecisional correlates of evidence accumulation might also be more readily observed when participants perform an error detection task (Murphy et al., 2015) or more frequently provide confidence ratings indicating an error had occurred (Feuerriegel et al., 2022), or when additional, decision-relevant stimuli are presented after the first-order decision (Grogan et al., 2023). This supports the notion of the Pe as a well-established correlate of error awareness (Falkenstein et al., 1991; O’Connell et al., 2007; Charles et al., 2013; Murphy et al., 2015), as well as the idea that a postdecisional ERP component can track the influence of additional stimuli after a decision has been made (Grogan et al., 2023). Taken together, these findings suggest that the link between Pe amplitudes and confidence depends on the extent to which postdecisional evidence accumulation is used to determine confidence judgments or error detection decisions, which may vary across decision-making tasks.

Our results also do not challenge the idea that, in some cases, decision reversals and confidence judgments can be substantively influenced by continued evidence accumulation, even without stimuli being presented between the choice and confidence rating. For example, changes of mind can be driven by stimuli that appear during (and immediately before) motor action execution, where these stimuli do not influence the first-order decision (e.g., Resulaj et al., 2009; Turner et al., 2022). However, we note that, in our dataset and most existing work involving difficult perceptual discrimination tasks, there is no clear evidence of covariation between Pe component amplitudes and confidence as would be expected from a substantial influence of postdecisional evidence accumulation.

### 4.3. Limitations

Our findings should be interpreted with the following caveats in mind. First, participants did not provide sufficient numbers of confidence ratings indicating that an error had occurred (i.e., maybe/probably/surely incorrect ratings). This is despite them making objectively erroneous decisions in a substantial proportion of trials (indicated by the accuracy plots in Figure 2A). This is consistent with distributions of confidence ratings in Ko et al. (2022) using an almost identical task, which was designed to elicit substantial variance in confidence ratings within each relative evidence condition. However, this meant that there were not enough trials to test for ERP correlates of certainty in having made an error. This may also be why we did not observe differences in Pe amplitudes across trials with correct responses and errors, as the Pe is more closely linked to error awareness rather than error commission (e.g., Charles et al., 2013; Murphy et al., 2015).

In addition, ranges of confidence ratings varied across individuals, which is why we used our linear regression approach to test for associations between *relatively* higher and lower confidence reports. This variability is likely to arise from different criteria being used for different confidence rating categories across participants (e.g., DeCarlo, 2010; Peters & Lau, 2015). In other words, one participant’s “probably correct” rating may not necessarily map onto the same internal degree of confidence as another’s “probably correct” rating, making comparison of individual ratings difficult at the group level. For this reason, we did not judge it meaningful to test for differences between specific pairs of confidence ratings (e.g., “probably correct” compared to “certainly correct”). This prohibited us from assessing non-linearities in the mapping of ERP amplitudes to confidence ratings. Small-N studies (Smith & Little, 2018) may be appropriate for better characterising the functional form of ERP component-confidence associations within individuals in contexts where confidence rating criteria are more stable over time.

We also note that our findings do not pertain to the validity of evidence accumulation models that are fit to patterns of task performance data but do not make predictions about neuroimaging measures (e.g., Ratcliff & Stams, 2009; Pleskac & Busemeyer, 2010). Our findings specifically relate to how such processes specified in these models might be implemented in the brain and reflected in measures of electrophysiological activity. It is possible that processes specified in these models are simply not indexed by the ERP components analysed in the lines of EEG studies mentioned here.

Lastly, even in analyses controlling for effects of relative evidence, the factor of absolute evidence was still varied across trials in order to produce variation in confidence ratings. It is possible that increasing overall brightness led to both increased confidence ratings (despite a drop in accuracy) and larger response-locked CPP amplitudes due to larger visual evoked potentials. As we replicated effects reported in Grogan et al. (2023), we do not believe that our results are simply a by-product of this confound. In addition, we did not observe associations between levels of absolute evidence and CPP amplitudes when holding confidence ratings constant. As absolute evidence manipulations necessarily entail changes in the overall intensity of stimulation, it may not be possible to completely control for this potential confound in future work.

### 4.4. Conclusion

We report evidence of a parietal ERP correlate of confidence during decision formation, which was not simply a by-product of changes in relative evidence or accuracy. However, we did not find a similar parietal correlate of confidence during the postdecisional time window that occurred between choice and confidence reports. Our findings reinforce the notion that processes occurring during decision formation are likely to substantively inform confidence judgments and should be considered in models of how we compute (and communicate) our degree of confidence in our decisions.

## Acknowledgements

This project was supported by an Australian Research Council Grant (ARC DP160103353) to S.B. and R.H. and an Australian Research Council Discovery Early Career Researcher Award to D.F. (ARC DE220101508). Y.H.K. was supported by a Julich - University of Melbourne Postgraduate Academy (JUMPA) scholarship. Funding sources had no role in study design, data collection, analysis or interpretation of results. The authors thank William Turner and Maja Brydevall for their support with study preparation.

## Notes

### Competing Interest Statement

The authors have declared no competing interest.

### Summary of Updates

Included analyses of effects of absolute evidence. Colour scheme of Figures 3 and 4 have been updated.

