## Supplementary Material for "Neural correlates of confidence during decision formation in a perceptual judgment task"

### Mixed Effects Regression Model Structures and Coefficients for Analyses of Accuracy, Response Times, and Confidence Ratings

Code for fitting all models listed below will be available at osf.io/7xv3m at the time of publication. All analyses were conducted in R (version 4.0.1). Generalised linear mixed effects models (GLMMs) were fitted using the lme4 package (version 1.1; Bates et al., 2015), statistical significance of each effect was determined by likelihood ratio tests conducted using the afex package (version 0.28; Singmann et al., 2017) in a stepwise forward approach, where each effect of interest was entered into the model and models before and after an effect was included were compared. Linear mixed-effects models with random intercepts for each participant were used to test the effects of relative evidence, absolute evidence, and their interaction on accuracy, response time (RT) and confidence rating measures.

#### List of Variable Names and Definitions

RT – The response time following the onset of the flickering squares in seconds

trial_outcome – The decision outcome following the flickering square (correct or error)

absolute_evidence (Abs) - The summed luminance of both squares

relative_evidence (Rel) – The difference in mean luminance across the brighter and darker squares

Luminance values in each condition varied with respect to the difference in mean luminance across the brighter and darker squares

confidence_rating – Decision confidence ratings between 1 – 7 (with 1 corresponding to surely incorrect, i.e., the minimum confidence rating, 4 corresponding to guessing/unsure, and 7 corresponding to surely correct, i.e., the maximum confidence rating)

ID – The participant ID number (typically used for defining random intercepts and slopes)

**Regression Model Equations and Coefficients**

Full model equations are listed below. Here, we follow the conventions of the lme4 package notation (Bates et al., 2015) when describing the structure of mixed effects regression models. Model coefficients for the full model (including the fixed effect of interest) are displayed below each set of model equations.

**Testing for the Effects of Relative Evidence, Absolute Evidence, and Their Interactions on Proportions of Correct Responses**

For analyses of proportion correct responses (i.e. accuracy), we fit generalised linear mixed effects regression models (Binomial family) with a logit link function.

trial_outcome ~ relative_evidence * absolute_evidence + (1 | ID)

Likelihood Ratio Tests Results for Predicting Accuracy (Log Odds of Being Correct) from Relative Evidence, Absolute Evidence, and Their Interactions

| Predictor | df | $\chi2$ | p |
|---|---|---|---|
| Rel | 2 | 746.23 | <.001*** |
| Abs | 2 | 225.62 | <.001*** |
| Rel × Abs | 4 | 26.62 | <.001*** |

Note. Rel: Relative evidence; Abs: Absolute evidence.
*p <.05 **p <.01 ***p <.001

Regression Coefficients for Predicting Accuracy (Log Odds of Being Correct) from Relative Evidence, Absolute Evidence, and Their Interactions

| Parameters | Estimate | SE | z | p |
|---|---|---|---|---|
| Intercept | 0.73 | 0.04 | 17.08 | <.001*** |
| Low Rel | -0.38 | 0.02 | -20.91 | <.001*** |
| Med Rel | -0.12 | 0.02 | -6.24 | <.001*** |
| Low Abs | 0.26 | 0.02 | 13.20 | <.001*** |
| Med Abs | -0.02 | 0.02 | -1.21 | .228 |
| Low Rel × Low Abs | -0.10 | 0.03 | -3.63 | <.001*** |
| Med Rel × Low Abs | -0.05 | 0.03 | -1.80 | .073 |
| Low Rel × Med Abs | 0.05 | 0.03 | 1.82 | .069 |
| Med Rel × Med Abs | 0.01 | 0.03 | 0.32 | .749 |

Note. Intercept represents the estimate for high relative and high absolute evidence.
Rel: Relative evidence; Abs: Absolute evidence.
*p <.05 **p <.01 ***p <.001

**Testing for the Effects of Relative Evidence, Absolute Evidence, and Their Interactions on RTs (Correct Response Trials)**

For analyses of RTs for correct trials, we fit generalised linear mixed effects regression models (Gamma family) with an identity link function.

RT ~ relative_evidence * absolute_evidence + (1 | ID)

Likelihood Ratio Tests Results for Predicting Response Time (Correct Trials) from Relative Evidence, Absolute Evidence, and Their Interactions

| Predictor | df | $\chi 2$ | p |
|---|---|---|---|
| Rel | 2 | 64.28 | <.001*** |
| Abs | 2 | 98.51 | <.001*** |
| Rel × Abs | 4 | 38.80 | <.001*** |

Note. Rel: Relative evidence; Abs: Absolute evidence.
*p <.05 **p <.01 ***p <.001

Regression Coefficients for Predicting Response Time (Correct Trials) from Relative Evidence, Absolute Evidence, and Their Interactions

| Parameters | Estimate | SE | z | p |
|---|---|---|---|---|
| Intercept | 790.47 | 2.92 | 270.88 | <.001*** |
| Low Rel | 8.07 | 1.70 | 4.73 | <.001*** |
| Med Rel | 8.91 | 1.64 | 5.42 | <.001*** |
| Low Abs | 21.16 | 1.82 | 11.65 | <.001*** |
| Med Abs | -5.40 | 1.99 | -2.72 | .007** |
| Low Rel × Low Abs | 13.26 | 2.10 | 6.33 | <.001*** |
| Med Rel × Low Abs | 4.62 | 2.46 | 1.88 | .060 |
| Low Rel × Med Abs | -7.21 | 2.13 | -3.38 | .001** |
| Med Rel × Med Abs | -3.81 | 2.08 | -1.83 | .067 |

Note. Intercept represents the estimate for high relative and high absolute evidence. Rel: Relative evidence; Abs: Absolute evidence.
*p <.05 **p <.01 ***p <.001

**Testing for the Effects of Relative Evidence, Absolute Evidence, and Their Interactions on RTs (Error Trials)**

For analyses of RTs for error trials, we fit generalised linear mixed effects regression models (Gamma family) with an identity link function.

RT ~ relative_evidence * absolute_evidence + (1 | ID)

Likelihood Ratio Tests Results for Predicting Response Time (Error Trials) from Relative Evidence, Absolute Evidence, and Their Interactions

| Predictor | df | $\chi 2$ | p |
|---|---|---|---|
| Rel | 2 | 3.21 | .201 |
| Abs | 2 | 28.92 | <.001*** |
| Rel × Abs | 4 | 14.08 | .007** |

Note. Rel: Relative evidence; Abs: Absolute evidence.
*p <.05 **p <.01 ***p <.001

Regression Coefficients for Predicting Response Time (Error Trials) from Relative Evidence, Absolute Evidence, and Their Interactions

| Parameters | Estimate | SE | z | p |
|---|---|---|---|---|
| Intercept | 814.86 | 3.19 | 255.27 | <.001*** |
| Low Rel | 3.19 | 2.43 | 1.31 | .190 |
| Med Rel | 3.67 | 2.51 | 1.46 | .144 |
| Low Abs | 19.47 | 2.15 | 9.07 | <.001*** |
| Med Abs | -13.11 | 2.16 | -6.07 | <.001*** |
| Low Rel × Low Abs | 15.34 | 2.84 | 5.40 | <.001*** |
| Med Rel × Low Abs | 2.35 | 3.12 | 0.75 | .451 |
| Low Rel × Med Abs | -2.93 | 2.71 | -1.08 | .280 |
| Med Rel × Med Abs | -4.86 | 3.00 | -1.62 | .106 |

Note. Intercept represents the estimate for high relative and high absolute evidence. Rel: Relative evidence; Abs: Absolute evidence.
*p <.05 **p <.01 ***p <.001

**Testing for the Effects of Relative Evidence, Absolute Evidence, and Their Interactions on Confidence Ratings (Correct Trials)**

For analyses of confidence ratings for trials with correct responses, we fit linear mixed effects regression models (Gaussian family).

confidence_rating ~ relative_evidence * absolute_evidence + (1 | ID)

Likelihood Ratio Tests Results for Predicting Confidence (Correct Trials) from Relative Evidence, Absolute Evidence, and Their Interactions

| Predictor | df | χ2 | p |
|---|---|---|---|
| Rel | 2 | 494.24 | <.001*** |
| Abs | 2 | 200.91 | <.001*** |
| Rel × Abs | 4 | 54.75 | <.001*** |

Note. Rel: Relative evidence; Abs: Absolute evidence.
*p <.05 **p <.01 ***p <.001

Regression Coefficients for Predicting Confidence (Correct Trials) from Relative Evidence, Absolute Evidence, and Their Interactions

| Parameters | Estimate | SE | z | p |
|---|---|---|---|---|
| Intercept | 5.77 | 0.11 | 52.91 | <.001*** |
| Low Rel | -0.21 | 0.01 | -17.82 | <.001*** |
| Med Rel | -0.02 | 0.01 | -1.43 | .154 |
| Low Abs | -0.15 | 0.01 | -13.17 | <.001*** |
| Med Abs | 0.02 | 0.01 | 1.50 | .132 |
| Low Rel × Low Abs | -0.10 | 0.02 | -6.02 | <.001*** |
| Med Rel × Low Abs | -0.00 | 0.02 | -0.09 | .931 |
| Low Rel × Med Abs | 0.03 | 0.02 | 1.82 | .068 |
| Med Rel × Med Abs | 0.00 | 0.02 | 0.26 | .795 |

Note. Intercept represents the estimate for high relative and high absolute evidence. Rel: Relative evidence; Abs: Absolute evidence.
*p <.05 **p <.01 ***p <.001

**Testing for the Effects of Relative Evidence, Absolute Evidence, and Their Interactions on Confidence Ratings (Error Trials)**

For analyses of confidence ratings for trials with error responses, we fit linear mixed effects regression models (Gaussian family).

confidence_rating ~ relative_evidence * absolute_evidence + (1 | ID)

Likelihood Ratio Tests Results for Predicting Confidence (Error Trials) from Relative Evidence, Absolute Evidence, and Their Interactions

| Predictor | df | $\chi 2$ | p |
|---|---|---|---|
| Rel | 2 | 128.07 | <.001*** |
| Abs | 2 | 239.46 | <.001*** |
| Rel × Abs | 4 | 3.95 | .413 |

Note. Rel: Relative evidence; Abs: Absolute evidence.
*p <.05 **p <.01 ***p <.001

Regression Coefficients for Predicting Confidence (Error Trials) from Relative Evidence, Absolute Evidence, and Their Interactions

| Parameters | Estimate | SE | z | p |
|---|---|---|---|---|
| Intercept | 5.10 | 0.10 | 51.71 | <.001*** |
| Low Rel | 0.20 | 0.02 | 9.38 | <.001*** |
| Med Rel | 0.06 | 0.02 | 2.82 | .005** |
| Low Abs | -0.36 | 0.02 | -15.14 | <.001*** |
| Med Abs | 0.12 | 0.02 | 5.46 | <.001*** |
| Low Rel × Low Abs | 0.01 | 0.03 | 0.32 | .751 |
| Med Rel × Low Abs | -0.04 | 0.03 | -1.25 | .212 |
| Low Rel × Med Abs | -0.02 | 0.03 | -0.60 | .549 |
| Med Rel × Med Abs | -0.01 | 0.03 | -0.27 | .791 |

Note. Intercept represents the estimate for high relative and high absolute evidence. Rel: Relative evidence; Abs: Absolute evidence.
*p <.05 **p <.01 ***p <.001

### Covariation Between Confidence, Accuracy and Response Times

We additionally examined how confidence ratings varied with accuracy and RT, in order to confirm that participants used the rating scale to meaningfully report their confidence. The overall mean accuracy (percent correct) was 66.53% (SE = 0.91%) and the mean response time was 784 ms (SE = 28ms). Supplementary Figure S1 displays accuracy (proportion correct) for each confidence level (Figure S1A), confidence rating distributions for correct and error trials (Figure S1B), and mean response times for each confidence level (Figure S1C).

Supplementary Figure S1A shows that accuracy was higher in trials with higher confidence ratings, indicating that participants reported confidence in a way that correlated with their objective decision accuracy. As visible in Supplementary Figure S1B, confidence ratings in the majority of trials (for both correct responses and errors) were between the range of "guessing" (4) to "surely correct" (7). There were relatively few trials with confidence ratings of "surely incorrect" (1), "probably incorrect" (2) or "maybe incorrect" (3). Supplementary Figure S1C shows that response times were negatively correlated with confidence (when confidence was > 4) for both trials with correct responses and errors.

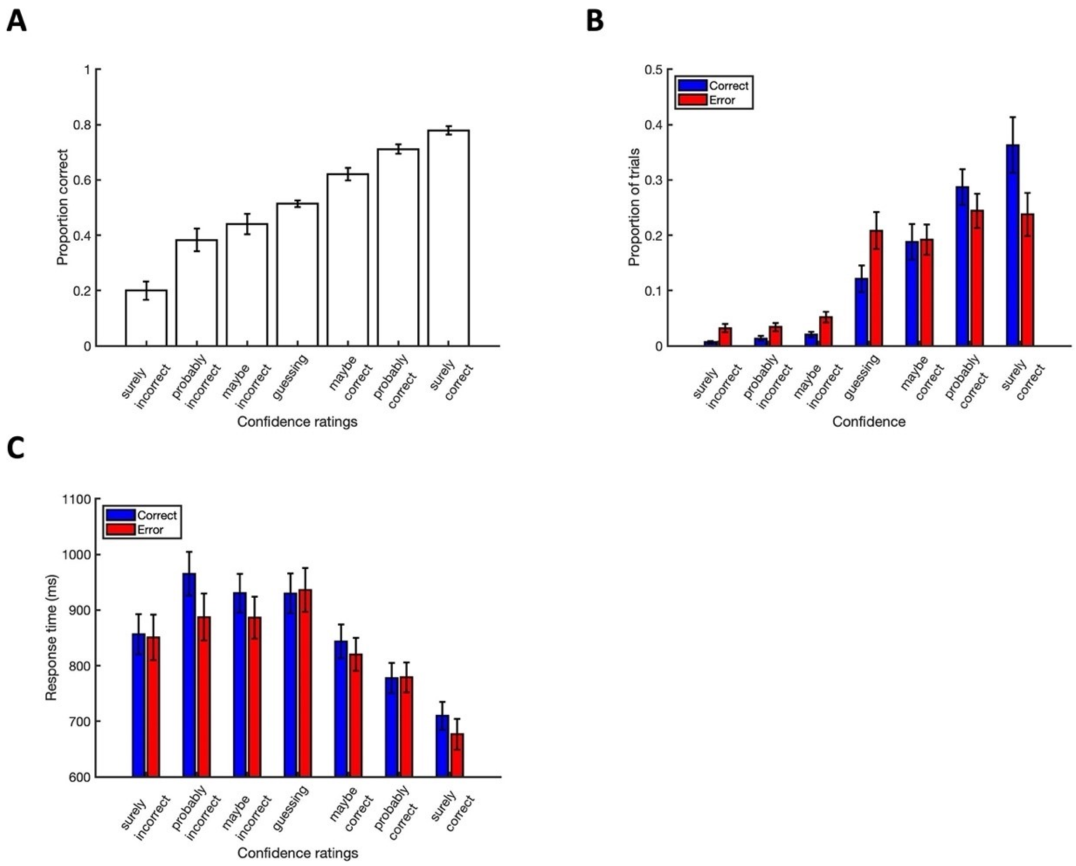


**Supplementary Figure S1.** Associations between accuracy, response times and confidence ratings. (A) Accuracy (proportion correct) for each confidence rating. (B) Confidence rating distributions for correct and error trials. (C) Mean RTs for each confidence rating for correct and error trials separately. Error bars represent SEM.

### Topographies of Associations between Confidence and CPP Amplitudes

To better visualise the topography of ERP-confidence associations during the pre-response CPP measurement window we performed regression analyses using confidence ratings to predict ERP mean amplitudes for each electrode separately, as done by Feuerriegel et al. (2022). Beta values (regression model slopes), intercepts and mean amplitude values were then averaged at the group level for each electrode and plotted. Scalp maps of group-averaged beta values, intercepts and mean amplitudes are visible in Supplementary Figure S2. Beta values (representing changes in amplitudes across confidence levels, Supplementary Figure 2A) were largest over parietal channels but were somewhat broad and left-lateralised. Intercepts (Figure S2B) did not show a clearly-defined topography, however mean amplitude values (Figure S2C) showed a topography that was typical of the scalp distribution of the CPP component (e.g., O'Connell et al., 2012).

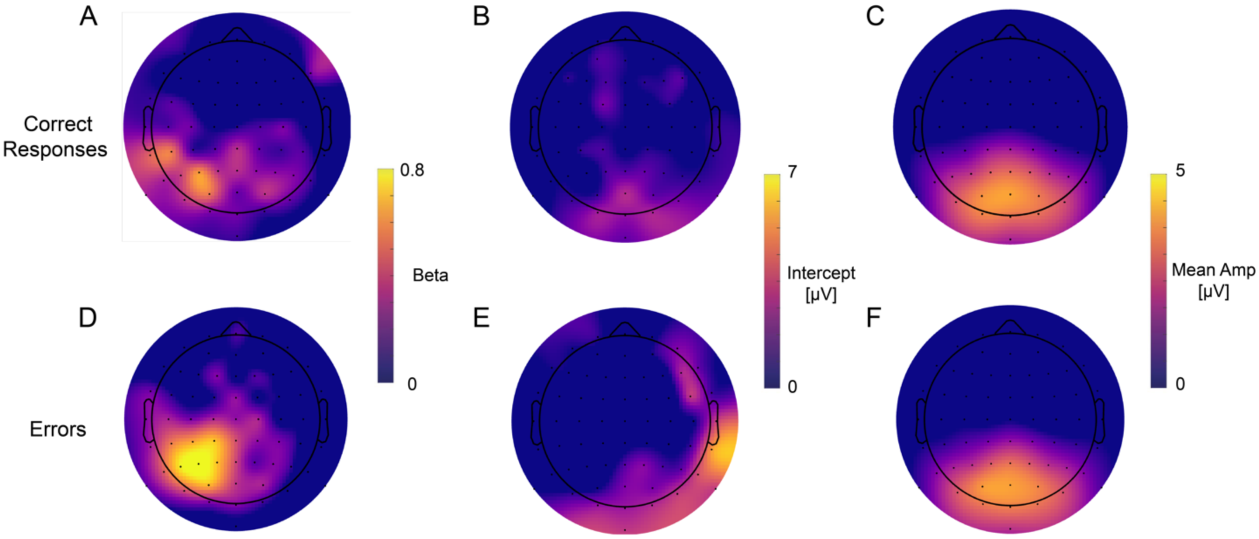


**Supplementary Figure S2.** Scalp maps of group-averaged beta values (A), intercepts (B) and mean amplitudes (C) from analyses predicting pre-response CPP amplitudes from confidence ratings ranging between "maybe correct" (5) and "surely correct" (7). Separate scalp maps are shown for trials with correct responses (top row) and errors (bottom row).

### Supplementary Material References

Bates, D., Maechler, M., Bolker, B., & Walker, S. (2015). Fitting linear mixed-effects models using lme4. *Journal of Statistical Software, 67*(1), 1-48.

Feuerriegel, D., Murphy, M., Konski, A., Mepani, V., Sun, J., Hester, R., & Bode, S. (2022). Electrophysiological correlates of confidence differ across correct and erroneous perceptual decisions. *NeuroImage, 259*, 119447.

O'Connell, R. G., Dockree, P. M., & Kelly, S. P. (2012). A supramodal accumulation-to-bound signal that determines perceptual decisions in humans. *Nature Neuroscience, 15*(12), 1729.

Singmann, H., Bolker, B., Westfall, J., Aust, F., & Ben-Shachar, M. S. (2017). afex: Analysis of factorial experiments. R package.
